# Water stable isotopes reveal the ecohydrological importance of stemflow for mature and juvenile European beech

**DOI:** 10.1101/2025.10.01.679712

**Authors:** Simon Keller, Laura Kinzinger, Judith Mach, Kathrin Kühnhammer, Markus Weiler, Natalie Orlowski, Christiane Werner, Simon Haberstroh

## Abstract

- Stemflow of forest trees can contribute a significant fraction of water to forest’s water fluxes; however, it is still unclear if and to what extent trees use stemflow for water supply and how much stemflow is lost by percolating below the root zone.
- We applied deuterium-enriched stemflow equivalent to a throughfall depth of 23 mm to adult *Fagus sylvatica* trees to trace stemflow through the soil and trees. We continuously measured *in-situ* water stable isotope compositions in soil and xylem water and destructively sampled xylem water in the crowns of adult labelled (n=18), unlabelled *F. sylvatica* (n=15) and unlabeled *Picea abies* (n=9), complemented by destructive xylem water sampling of neighboring juvenile *F. sylvatica* (n=45).
- Stemflow water supported 3.9 – 14.0% of daily sap flux of adult labelled *F. sylvatica* trees. In the soil, deuterium-enriched stemflow was detectable at a max. of 0 – 0.40 m to the labelled tree. However, unlabeled juvenile trees within a distance of ∼2 m showed label water uptake, indicating rooting into soil compartments affected by stemflow label.
- We demonstrate the importance of stemflow as a water source for both adult and neighboring juvenile *F. sylvatica*, strongly profiting from stemflow infiltration.

## Introduction

In forested ecosystems, the vegetation actively divides the precipitation input into several fractions, i.e., interception, throughfall and stemflow. The latter denotes the precipitation, which drains from leaves and branches to the stem, where it is channelled down the trunk (Levia *et al*., 2011). Stemflow is a highly variable phenomenon, depending on meteorological conditions, stand density and tree species, which determines the surface texture of the bark and the shape of the plant (Levia & Germer, 2015). Therefore, the share of stemflow in relation to total precipitation varies greatly. For European Beech (*Fagus sylvatica* L.), a species with smooth bark, proportions between 5 and 8% of total precipitation have been reported in Central Europe (Staelens *et al*., 2008; Levia *et al*., 2010). Although this may appear to be a small share, the concentration of stemflow within local areas at the base of the trunk has potentially large impact on the magnitude and timing of water inputs into the soil (Carlyle-Moses *et al*., 2018; Tischer *et al*., 2020; Metzger *et al*., 2021). Stemflow redistributes and retains precipitation input, creating spatiotemporal heterogeneity in soil surface moisture close to the tree stem (Tischer *et al*., 2020). Particularly in species with higher stemflow proportions, such as *F. sylvatica*, stemflow can lead to increased root-induced bypass flow (Schwärzel *et al*., 2012; Pinos *et al*., 2023). The “double-funnelling of trees”, introduced by Johnson & Lehmann (2006) refers to this two-sided process, where on the one hand, aboveground rainfall is channelled and concentrated along the stem, and on the other hand, sub-soil preferential flow occurs along roots, which can have both, vertical and lateral effects.

Recently, there has been growing interest in stemflow, particularly in respect to the infiltration area (the soil surface area the stemflow water spreads to during infiltration) (van Stan & Allen, 2020; Metzger *et al*., 2021; Llorens *et al*., 2022). While infiltration areas of up to 11.8 m² have been reported (van Stan & Allen, 2020), including species from different biomes, such as temperate, tropical and Mediterranean, typically the infiltration area of stemflow for average precipitation events has been found to be rather small with < 1 m² (Carlyle-Moses *et al*., 2020; Llorens *et al*., 2022; Zuecco *et al*., 2025). It remains uncertain if and to what proportion trees use stemflow funnelled water for their own water fluxes (Snyder *et al*., 2024; van Stan & Pinos, 2024). As stemflow can increase soil water resources near the stem (Hemr *et al*., 2023), other individuals of the same or different species might also profit from these water inputs, e.g., by rooting horizontally into wetter soil compartments, competing for these water resources with the trees that produced the stemflow (van Stan & Pinos, 2024). While several studies suggest that, especially in semi-arid climates, stemflow is a readily accessible water resource for water uptake in the rooting zone of plants (Li *et al*., 2009; Wang *et al*., 2019; Snyder *et al*., 2024), experimental studies on this topic are scarce and our quantitative knowledge remains limited. In previous studies, various methods were applied to investigate stemflow, primarily the analysis of soil chemical properties or infiltration experiments with dyes (Metzger *et al*., 2021; Llorens *et al*., 2022). While these methods primarily investigate stemflow infiltration, it is not possible to trace the path of stemflow water through the vegetation. Recently, stemflow was investigated using the ratio of stable isotopes of hydrogen (δ^2^H) and oxygen (δ^18^O) (Pinos *et al*., 2023; Snyder *et al*., 2024; Zuecco *et al*., 2025). This approach has not yet been widely used to characterize stemflow and its implications for tree water fluxes. So far, only Snyder *et al*. (2024) have estimated the proportion of stemflow in the xylem water of two dryland species (*Pinus monophylla* and *Juniperus osteosperma*) in the Great Basin (USA). Their results indicate contributions of stemflow of 0-2% to total xylem water of the two species. However, they used a rather low label strength (δ^2^H = 200 - 400 ‰) and a low amount of water for stemflow (1.1 - 17 L) without mimicking co-occurring precipitation, potentially resulting in a high noise-to-signal ratios (Snyder *et al*., 2024).

Water stable isotopes are an established tool in ecohydrological research to assess potential sources of root water uptake by comparing xylem water isotopic signatures with those of available water (Ehleringer & Dawson, 1992), such as soil water across depth. To investigate specific water fluxes, labelling with ^2^H or ^18^O enriched water (Werner *et al*., 2021; Kinzinger *et al*., 2025) provides an alternative to other tracer methods (e.g. with dyes, van Stan & Allen (2020); Pinos *et al*., (2022)), having the advantage that both, infiltration area (Zuecco *et al*., 2025) and contribution of stemflow to tree water fluxes (Snyder *et al*., 2024) can be estimated. In addition, *in-situ* water isotope probes allow the direct and continuous measurement of water stable isotope ratios in the soil and xylem (Volkmann & Weiler, 2014; Seeger & Weiler, 2021; Kühnhammer *et al*., 2022; Kinzinger *et al*., 2024). These *in-situ* water isotope measurements drastically expand our possibilities to investigate temporal dynamics in ecosystem water fluxes. However, to simultaneously achieve a high spatial resolution, traditional methods, such as destructive sampling for cryogenic vacuum extraction, remain viable and valuable additions (Haberstroh *et al*., 2024). Furthermore, cryogenic vacuum extraction is more cost efficient compared to new *in-situ* systems, especially with a large number of samples (Fischer *et al*., 2019). Thus, the potential of water stable isotopic labelling approaches to elucidate the spatial and temporal impact of stemflow water on ecohydrological processes, such as tree water fluxes, is extremely high.

Here, we trace stemflow with deuterium-enriched water in mature *F. sylvatica* trees, while applying the corresponding amount of a natural throughfall event as non-labelled water. We combine isotopic labelling approaches with a novel *in-situ* system and sap flow measurements. This enables us continuously to qualitatively and quantitatively trace the uptake of stemflow water by the tree and its surrounding trees in daily resolution.

We hypothesize that stemflow of mature *F. sylvatica* is a readily available and relevant water source for adult trees generating stemflow, but also for understory vegetation and close neighbouring trees. This indicates that the spatial distribution of stemflow in the vegetation extends beyond local infiltration hotspots, i.e., in particular younger neighbouring trees can reach stemflow infiltration areas of the older trees with their roots and make use of these water resources.

## Material and Methods

We conducted our experiment from July 1 to August 31 2023 in a temperate mixed forest, dominated by mature European beech *(Fagus sylvatica* L.) and Norway spruce *(Picea abies* L.), interspersed with individual specimens of *Quercus petraea* L., *Pseudotsuga menziesii*, *Abies alba* and other tree species (Kinzinger *et al*., 2024; Dumberger *et al*., 2025). The experimental plot is situated on the western slope of the Black Forest, Ettenheim, Germany (48.254417, 7.924278 UTM WSG84, 530 - 540 m a.s.l.) (Kinzinger *et al*., 2024) within the ECOSENSE forest (Werner *et al*., 2024). The site with an area of 0.75 ha is located near the hilltop of a west facing slope (6°-10°). Mean annual air temperature and mean annual precipitation sum are 10.4°C and 1070 mm, respectively (DWD, 2022). The soil with a depth of 0.90 – 1.00 m is classified as Cambisol (LGRB 2021). The understorey is dominated by juvenile *F. sylvatica* trees. Local meteorological data, i.e., air temperature and relative humidity (CS215-L, Campbell Scientific Ltd., Bremen, Germany), precipitation (tipping bucket, 0.2 mm resolution, Davis Instruments, Hayward, USA) and photosynthetic active radiation (PAR) (LI-1500, LI-COR Biosciences GmbH, Bad Homburg, Germany) were measured at an open site approx. 600 m northwest of the study site. Data was obtained at an interval of five minutes and stored on a data logger (CR-1000, Campbell Scientific, Shepshed, UK).

### Experimental design

The labelling experiment was carried out on six individual experimental plots distributed across the area. Three plots (pure stands, 160 m² each), contained solely *F. sylvatica*. The other three plots (320 m² each), represented mixed stands with approx. 50% *F. sylvatica* and 50% *P. abies* adult trees (Fig. 1) (Kinzinger *et al*., 2024). All plots contained at least three adult trees (between 60 and 80 years, 24-31 m height), and between 6-10 juvenile *F. sylvatica* trees. Adult *F. sylvatica* trees had an average diameter at breast height (DBH) of 31.8 ± 8.9 cm (n = 36, ± 1SD); juvenile *F. sylvatica* trees were smaller with a DBH of 8.7 ± 2.9 cm (n = 45) and a height between 2-8 m. *P. abies* trees on the label plots had a similar DBH (38.2 ± 5.9 cm, n = 9). For comparison, three *P. abies* plots, which were not directly involved in the experiment were included as control trees (n = 9), with a DBH of 38.0 ± 5.6 cm (n = 9). Volumetric soil water content was measured in four depths (0.05, 0.20, 0.40, 0.90 m) with SMT100 sensors (Truebner GmbH, Neustadt, Germany) in each irrigated experimental plot (n = 6) and recorded in 10-minute intervals.

**Figure 1:**
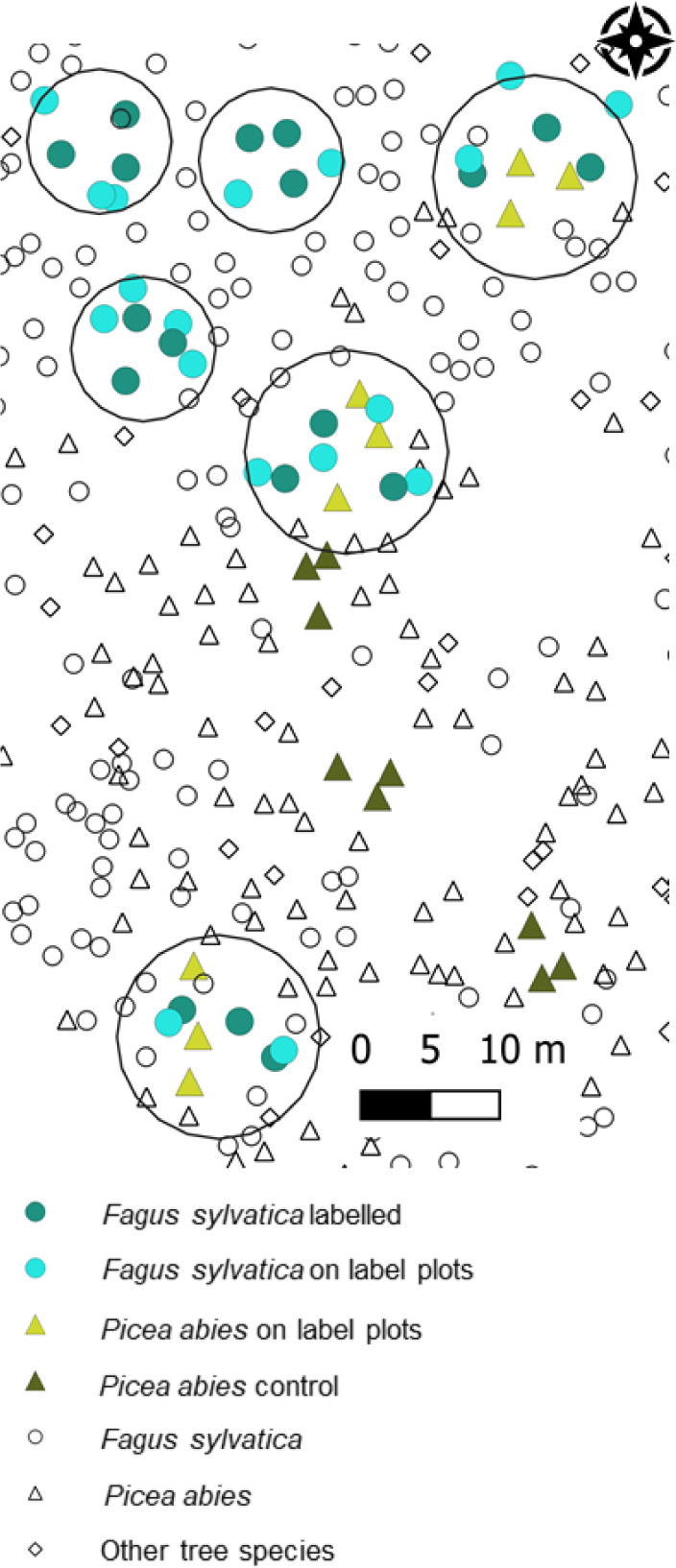
Field site and experimental plots including *F. sylvatica* trees with labelled stemflow (n = 18), unlabelled *F. sylvatica* on label plots (n =18), *P. abies* on label plots (n = 9) and *P. abies* as control trees (n = 9). The circles correspond to the plot sizes of 160 m² and 320 m², respectively, dependent on the presence of *P. abies* (Kinzinger *et al*., 2024). Juvenile *F. sylvatica* trees are not displayed.

On July 20, the amount of water corresponding to a precipitation event with approx. 23 mm throughfall was applied evenly with garden sprinklers on all six plots, leading to a precipitation duration of 1.5 hours per plots in pure stand and 3 hours per plot in mixed stand, corresponding to the respective area. The irrigation water had an isotopic signature of –58.0 ± 0.7‰ and – 8.70 ± 0.19‰ for δ^2^H and δ^18^O, respectively.

Stemflow was simultaneously applied on three adult beech individuals per plot (in total 18 of 36 trees), using deuterium-enriched water with mean δ^2^H and δ^18^O values of +5,822 ± 214‰ and –6.84 ± 0.16‰, respectively. Stemflow amount was adjusted for each individual by calculating the average funnelling ratio (*FR*) of *F. sylvatica* trees on site for the applied event of 23 mm (see equation 1) using data obtained by stemflow measurements on site (n = 6; see (Kinzinger *et al*., 2024)):

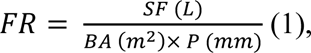

Where *SF* is stemflow, *BA* basal area and *P* precipitation. Subsequently, we determined stemflow volume (*SV*) for each investigated tree *i* (equation 2):

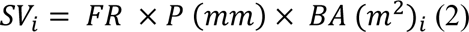

For stemflow application, hoses with holes were placed horizontally around the stem at around 0.75 m height and attached to a canister (Fig. S1). Flow rates were adjusted by the number of holes, aiming for a consistent flow duration of three to four hours per tree, simulating a small retention compared to the precipitation, which is typical for stemflow (Carlyle-Moses *et al*., 2018). Although stemflow sometimes naturally leads to irregular infiltration patterns into the soil, we aimed for a relatively even circumferential distribution of the applied water to avoid biases due to unilateral infiltration.

### *In-situ* **water stable isotope measurements**

*In-situ* isotopic values of soil and xylem water were measured according to the “diffusion-dilution sampling” method by Volkmann & Weiler (2014), using an automated soil and water isotopic monitoring system (Seeger & Weiler, 2021; Kinzinger *et al*., 2024; Kinzinger *et al*., 2025). Microporous probes were installed in four soil depths (0.05, 0.20, 0.40, 0.90 m) at the centre of each plot and at the base of stems of mature *F. sylvatica* (n = 33, n = 18 with labelled stemflow, n = 15 without label) and *P. abies* trees (n =18, n = 9 in stemflow plots, n = 9 in non-stemflow plots). Tubes were connected directly to a Cavity Ring-Down Spectroscopy (CRDS) water isotope analyser (L2130*-i*, Picarro Inc., Santa Clara, CA, USA), using fluorinated ethylene propylene tubing (see experimental setup by Kinzinger *et al*. (2024). Three water isotope standards (see Table S1), calibrated against VSMOW-2, SLAP-2 and GRESP (IAEA), stored in airtight containers and buried below the soil surface, were automatically measured every 1-2 days. An average of 120 sec. was calculated for isotopic measurements, when values reached a stable state, defined, when the following conditions were met over the last 150 sec.: H_2_O_sd < 150.0, δ^18^O_sd < 0.30, δ^2^H_sd < 0.90, H_2_O_ti < 150, δ^18^O_ti < 0.05, δ^2^H_ti < 0.30 (sd: standard deviation, ti: trend index) (for details see Seeger & Weiler, 2021). There was a linear relationship between water vapor concentration (ppmV) and δ^2^H/ δ^18^O measurements for the standards, therefore all measured values were normalized to a medium ppmV value of 13,500 according to this relationship. Afterwards, we converted measured vapor values to liquid isotopic composition using a linear regression between liquid and vapor measurements established for the standards (Kinzinger *et al*., 2024). *In-situ* isotopic data was carefully checked to exclude values, which were unrealistic caused by errors in the measurement system. This included, for example, δ^18^O values, which were more enriched than all precipitation or soil measurements of the year 2023, as this pointed to condensation effects in the measuring lines.

### Destructive sampling and measurements

We collected destructive plant samples in addition to the continuous *in-situ* isotopic measurements for all trees on the label plots (Fig. 1), starting with a baseline sampling two days before labelling (see Table 1 for timing of destructive sampling). For juvenile trees (n = 45), samples were always taken at 2-3 m height, independent of the tree height. Samples from all mature *F. sylvatica* (n = 36, 18 labelled, 18 unlabelled) and *P. abies* (n = 9) trees on the experimental plots were sampled from the sun crown with the help of tree climbers. Due to weather conditions, not all adult trees could be sampled every time, however, we were always able to obtain at least nine replica per group of mature trees (as defined in Table 1). To obtain xylem water samples and following guidelines by Ceperley *et al*. (2024), phloem was removed from all branches as it can affect the water isotope signature due to contained organic substances. The branches were cut into small pieces, stored in 12 ml vials (Labco® Exetainer, Labco Limited, Lampeter, UK) and lids were additionally sealed with Parafilm.

**Table 1:** Sampling frequency for soil (n = 6 per depth) and tree crown xylem water for the different sampling groups.

| Group/Date | 18/07 | 24/07 | 26/07 | 27/07 | 31/07 | 2/08 | 8/08 | 15/08 |
| --- | --- | --- | --- | --- | --- | --- | --- | --- |
| **Juvenile *F. sylvatica***<br>**(n = 45)** | X | X |  | X | X |  | X | X |
| **Stemflow labeled**<br>***F. sylvatica***<br>**(n = 18)** | X |  | X |  |  | X | X | X |
| **Non- labeled**<br>***F. sylvatica***<br>**(n = 18)** | X |  | X |  |  | X | X | X |
| ***P. abies* (n = 9)** | X |  | X |  |  | X | X | X |
| **Soil (n = 6)** | X | X |  |  |  | X |  | X |

Note: On July 20, the labeling experiment was conducted.

In addition, soil samples were taken from each plot (sample frequency see Table 1). For this, one out of the three stemflow labelling trees per plot (n = 6) was chosen randomly, and soil samples were taken at three distances (0.10 m, 0.40 m and 1.00 m) on the south side of this tree. At 0.40 m and 1.00 m distance, three depths, i.e., 0.05 m, 0.20 m and 0.40 m, were sampled using a soil corer (length 1 m, core diameter 2.8 cm), resulting in six replica per distance, depth and sampling point. At 0.10 m, we only sampled soil in 0.05 m depth due to the extensive rooting system of *F. sylvatica* close to the stem. Additionally, on the evening following the labelling experiment, soil samples were taken in immediate proximity to six stemflow labelling trees in order to determine the label strength in the soil shortly after label application.

All xylem and soil samples were kept cool in the field. Due to the large number of samples, we could not ensure that all samples were extracted in time (2-8 weeks according to Ceperley *et al*. (2024)), thus we decided to freeze them at −20°C until further analysis (Dubbert *et al*., 2019). Plant and soil water were extracted using a custom-made cryogenic vacuum extraction line (Kübert *et al*., 2020) following the procedure detailed in Haberstroh *et al*. (2024). No cover material was added to plant samples; glass wool (extra fine, Karl Hecht GmbH & Co KG) was used for soil samples (Haberstroh *et al*., 2024). Shortly, under vacuum conditions (max 0.1 mbar, XDS10 vacuum pump, Edwards), samples were heated to 95 °C in a water bath for at least 90 minutes. The water evaporating from the samples froze out in connected glass U-tubes immersed in liquid nitrogen (–196 °C) (Kübert *et al*., 2020). After defrosting, samples were filtered using a glass fibre syringe filter (0.7 µm, Syringe Filter; Membrane Solutions) and stored at 6°C until further processing. Between 500 and 1500 µl of water was extracted per sample, which significantly minimizes the risk of ^2^H fractionation by organics compared to lower extracted amounts (Diao *et al*., 2022). The quality of the extraction was determined by weighing some of the samples after cryogenic vacuum extraction and again after drying them at 60 °C (to minimize carbon losses from plant materials) for 48 hours (Millar *et al*., 2022).

Stable isotopic signatures of the extracted xylem and soil water and of the precipitation samples were measured using CRDS (L2130-*i*, Picarro Inc., Santa Clara, CA, USA). To minimize memory effects, every xylem and soil water sample was measured nine times (Haberstroh *et al*., 2024). Samples which contained a significant enrichment in ^2^H, were measured 12 times. Precipitation samples were measured six times. The last three measured values were used for analysis if the δ^2^H values were within a range of ± 0.5‰ and δ^18^O values within a range of ± 0.1‰. Otherwise, the sample was measured again. Care was taken to ensure that the water vapor mixing ratio was around 17,000 ppmV for all samples, ensuring comparability of values (Haberstroh *et al*., 2024). We acknowledge that organic contaminations of samples can interfere with CRDS measurements of δ^2^H and δ^18^O (Hendry *et al*., 2011; Herbstritt *et al*., 2024). However, as we applied label with extremely high concentrations of ^2^H (δ^2^H = 5,822 ± 214‰), we expect the impact of organics on identifying label in xylem and soil water to be negligible. In addition, we deployed an *in-situ* measurement system for comparison, which has been found to reliably measure δ^2^H and δ^18^O (Beyer *et al*., 2020; Herbstritt *et al*., 2024). Additionally, we collected precipitation samples in three throughfall collectors distributed among the different plots on the field site. Samples were taken a total of eight times during the study period.

Isotopic composition of the extracted xylem and soil water, as well as precipitation water was calibrated using in-house standards regularly calibrated against the international standards VSMOW-2, SLAP-2 and GRESP (IAEA). For isotopic values of all used standards we refer to Table S1.

### Sap flow measurements

Sap flow was measured with heat pulse velocity sensors (HPV-06, Implexx, Melbourne, Australia) on mature *F. sylvatica* (n = 18) and *P. abies* (n = 18) trees in 10 mm and 20 mm xylem depth. Sensors were installed at 1.3 m height. Measurements were conducted every 10 min and data stored on a CR1000 datalogger (Campbell Scientific Ltd, Shepshed, UK). Sap flow was calculated following the Dual Method Approach (Forster, 2019, 2020), which is based on the heat ratio method (HRM; (Burgess *et al*., 2001)) and the Tmax method (Cohen *et al*., 1981), overcoming the limitations the methods possess at high and low flow rates, respectively (Forster, 2019). Probe misalignment was corrected according to Kinzinger *et al*. (2024) and Dumberger *et al*. (2025). To scale sap flow to total tree water fluxes (in L), the active sap wood area was determined by taking tree cores at the sensor height and using the light transmission method (Munster-Swendsen, 1987; Børja *et al*., 2016; Quiñonez-Piñón & Valeo, 2018; Kinzinger *et al*., 2024).

### Statistical analysis and Bayesian mixing model

To test the spatial distribution of the stemflow label, we analysed the relationship between the δ^2^H values of soil and xylem water samples (for soil and juvenile *F. sylvatica*) and the distance to the closest labelled *F. sylvatica* tree. Here, we used a three-parameter exponential decay function (*drm* function, R package *drc* (Ritz *et al.,* 2015)). This function showed the best fit to the data. This analysis was done separately for each sampling day, where a relationship between δ^2^H values and distance was found with the *drm* function.

Bayesian mixing models, such as *MixSIAR* (Stock *et al*., 2018) in R (R Core Team, 2023), are commonly used to calculate the share of different water sources to plant water uptake (Gessler *et al*., 2022; Kinzinger *et al*., 2024; Sprenger *et al*., 2025). As input data, we used δ^2^H and δ^18^O values of xylem water (at the stem base or in the tree crown) as ‘consumer’ data and δ^2^H and δ^18^O values of soil water in three different depths (0.05. 0.20 and 0.40 m, measured with the *in-situ* system), precipitation water, irrigation and stemflow water as ‘source’ data. As we were not able to measure water stable isotopes in the soil every day, the last available values for δ^2^H and δ^18^O were used in the model until a new measurement became available. Water source contribution by *MixSIAR* was estimated separately for xylem water at the stem base of labelled *F. sylvatica* trees, for xylem water in the tree crown of labelled *F. sylvatica* trees and in the tree crowns of juvenile *F. sylvatica* trees. In all models, no informative prior was formulated, as priors can have a strong impact on model outputs, potentially introducing biases (Stock *et al*., 2018). In addition, we did not introduce any concentration dependencies, as soil water availability was sufficient for root water uptake in all measured soil layers. The models were run with the Markov chain Monte Carlo function, using runtimes from “normal” to “very long”, aiming for a Gelman-Rubin value below 1.05 for all chains, considering a residual process error structure. For all modeling results including uncertainties see Table S2. The daily contribution of stemflow to tree water fluxes was calculated by multiplying the relative share of stemflow water determined by *MixSIAR* with total tree water fluxes in L day^−1^, considering error propagation. All data was analysed using R (version 4.3.2) (R Core Team, 2023).

## Results

### Meteorological conditions and sap flow

During the experimental period from July 1 to August 31, a mean air temperature of 19.5 °C was recorded with a precipitation sum of 262 mm (Fig. 2A), which was slightly wetter than in preceding years. A remarkably large precipitation event of 40.6 mm occurred on July 25, five days after the stemflow labelling (July 20). The following days remained cool and wet with frequent precipitation events. From August 7 onwards, mean air temperature increased. A sharp decline in air temperature to a minimum of 12.5°C (daily average), accompanied by heavy precipitation events occurred at the end of the study period. Prior to the stemflow labelling (1 July – 20 July), soil water content in the top three soil depths (0.05, 0.20, 0.40 m) was rather low with 11.0 ± 0.9, 10.1 ± 1.1 and 12.7 ± 1.0 %, respectively, while soil water resources in 0.90 m were less depleted (15.4 ± 0.7 %, Fig. 2B). Afterwards, soil moisture increased rapidly in all depths in response to the large precipitation event on July 25. Interestingly, we only detected a small increase in soil moisture in 0.20 and 0.40 m following the experimental irrigation on July 20 (Fig. 2B). Daily sap flow for mature *F. sylvatica* trees showed maximum values of 64.5 ± 10.5 L day^−1^ (9 July, Fig. 2C) in the study period. *P. abies* trees showed lower sap flow with maximum rates at 15.0 ± 3.9 L day^−1^ (18 August). In general, sap flow rates aligned well with meteorological conditions, such as air temperature and occurrence of precipitation, for both species (Fig. 2A,C).

**Figure 2:**
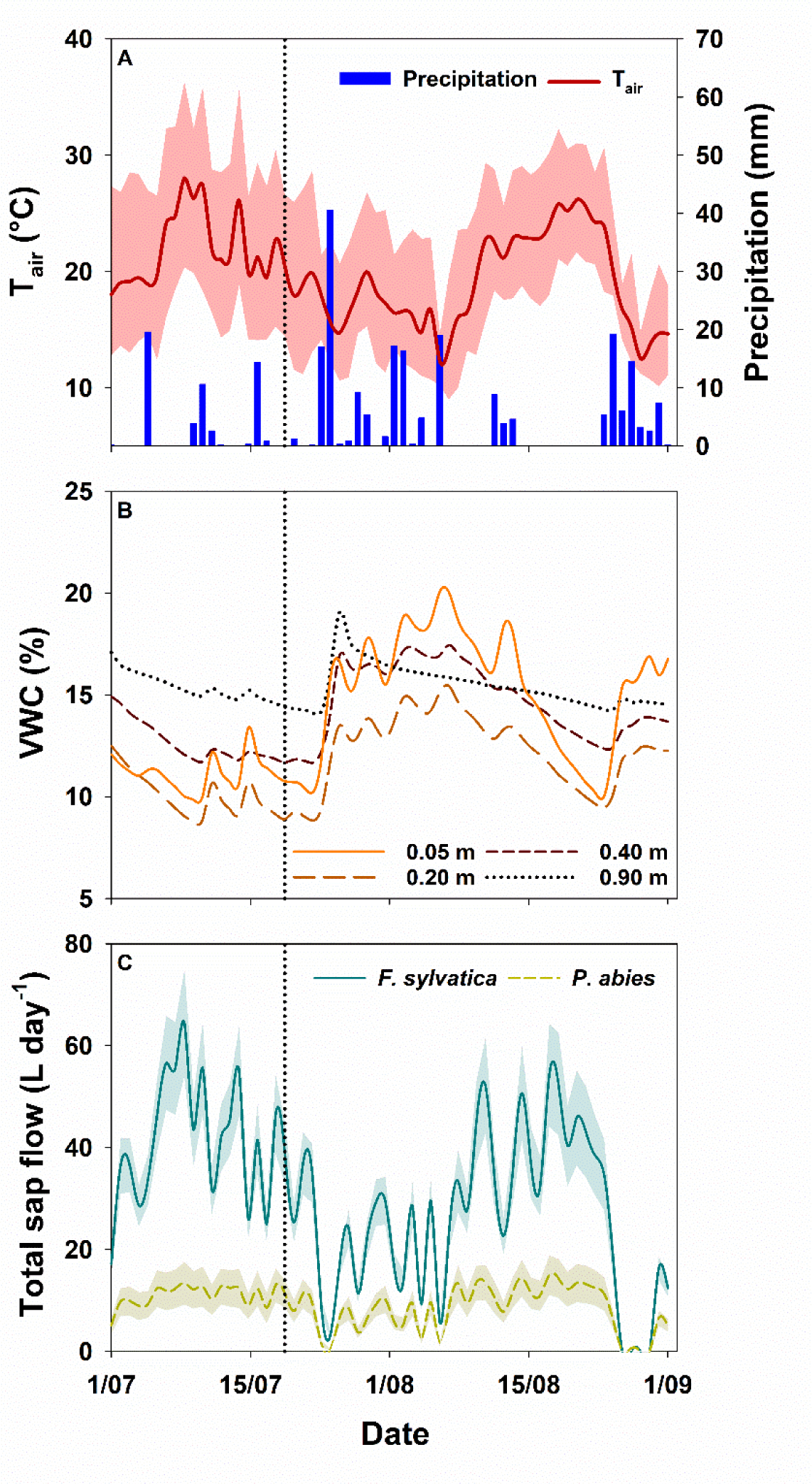
Meteorological conditions and mean daily sap flow during the experiment. Daily mean air temperature with minimum and maximum values (A), daily precipitation sum, daily mean soil volumetric water content (VWC) for different depths averaged over all six irrigated plots (n = 6 per depth) (B) and daily sap flow for examined *F. sylvatica* (n = 18) and *Picea abies* (n = 18) trees ± 1 SE. The dotted line marks the day of stemflow labelling and irrigation.

### Temporal dynamics of water stable isotopes in soil, xylem and tree crown water

We labelled stemflow of mature *F. sylvatica* trees during a simulated throughfall event of 23 mm and continuously followed its pathway with water stable isotopes in soil and xylem water at the stem base, as well as in the tree crowns with point measurements.

Measurements of δ^2^H in soil water revealed a rather localized effect of the stemflow labelling (Fig. 3A, B). In a distance of 0.10 m to the stem of labelled trees, large enrichment of δ^2^H = +1,745.8 ± 827.3 ‰ was found close to the soil surface (just below the litter) on the evening of the experiment (Fig. 3A). However, δ^2^H values declined strongly in the following days to δ^2^H = –25.2 ± 12.8 ‰ on August 2, which was within the range of recently fallen precipitation (Fig. 3B). Soil water at further distances to the stem showed no traces of the applied label with one exception in 0.40 m, where δ^2^H values increased from –26.2 ± 3.6 ‰ (pre-label) to –14.7 ± 21.9 ‰ on August 2. However, this occurred only in one single sample (δ^2^H = +298.1 ‰). Soil water samples at 1.00 m and further away from the stem mostly resembled dynamics of the precipitation natural isotopic composition and ranged from δ^2^H = –29.4 ± 14.0 ‰ to δ^2^H = –58.1 ± 7.0 ‰ (Fig. 3B). No significant differences between the sampled soil depths could be detected, thus averages are displayed in Fig. 3B. In addition, the resemblance of samples extracted with cryogenic vacuum extraction (distance 1.00 m) and measured with the *in-situ* system (distance > 1.00 m) indicate a general comparability between both approaches.

**Figure 3:**
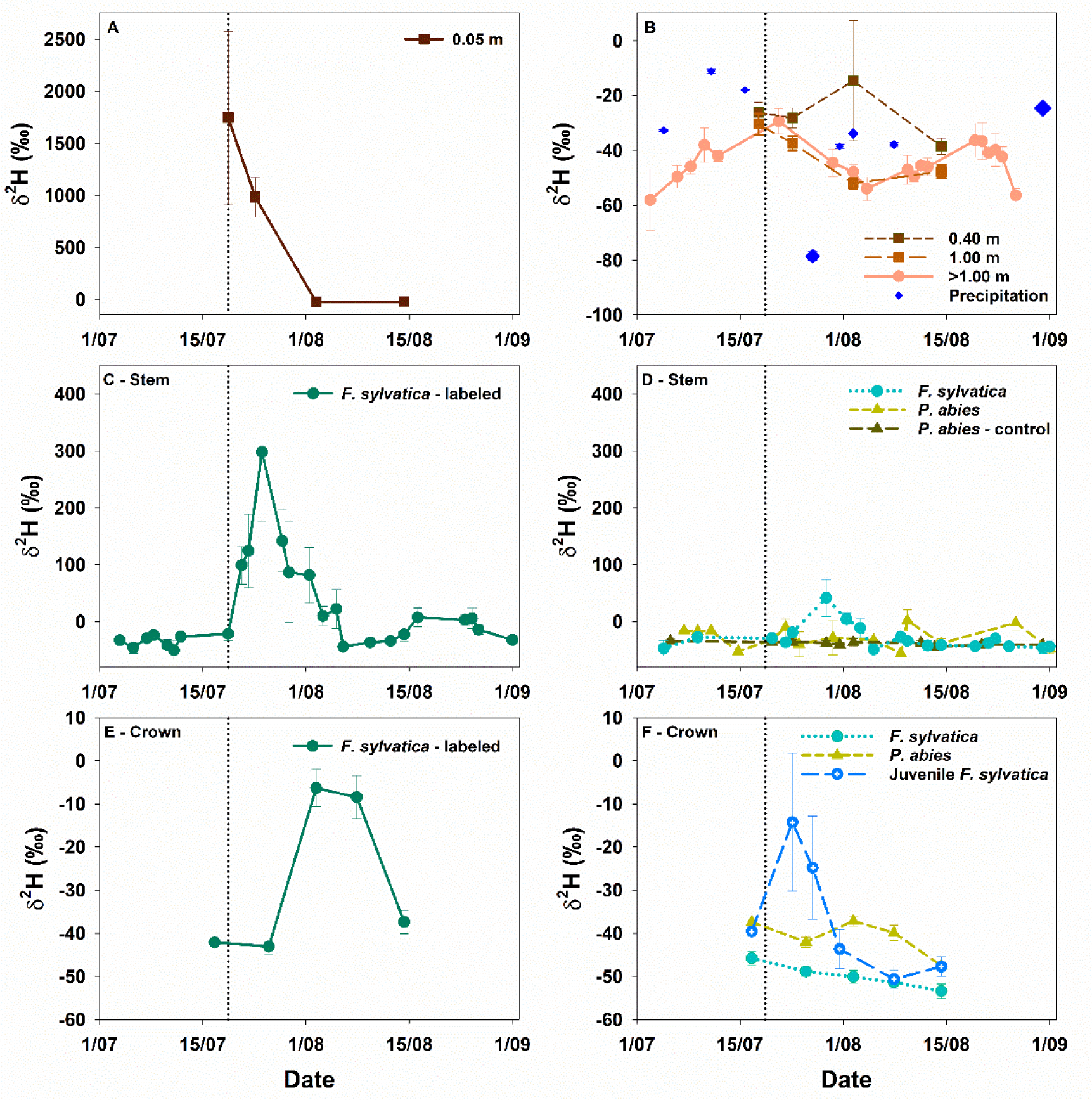
Temporal dynamics of measured δ^2^H values in soil water with standard error for soil water measured at 0.10 m distance to the labelled tree (n = 6) (A), for precipitation (n = 3, size corresponds to size of precipitation event) and soil water measured at different distances to the labelled tree (0.40, 1.00 and > 1.00 m) (n = 6 profiles) (B), xylem water of labelled *F. sylvatica* trees at the stem base (n = 3-11 per day) (C) and xylem water of unlabelled *F. sylvatica* (n = 3-9 per day) and *P. abies* trees (n = 2-5 per day), including control trees for *P. abies* (n = 3-6 per day), at the stem base (D). δ^2^H values of xylem water in the tree crown of labelled *F. sylvatica* trees (n = 18) (E) and xylem water in the tree crown of unlabelled *F. sylvatica* (mature (n = 19) and juvenile (n = 45)) and *P. abies* trees (n = 9) (F). All measurements are displayed with ±1 SE. The dotted line marks the day of stemflow labelling and forest irrigation. Precipitation was always sampled on the date shown and can represent a mixture of several small events. Note the different y-axis in panel A and B.

Similar to the enrichment in soil water close to the stem (Fig. 3A), we found increasing δ^2^H values in the stem xylem water of labelled trees already two days after label application (Fig. 3C). Already three days later (July 25), the maximum δ^2^H value (+298.0 ± 122.2 ‰) was reached and until August 6 (i.e. 17 days after label application) δ^2^H values dropped to pre-label values (Fig. 3C), only to increase again in mid-August. This secondary enrichment corresponded to the occurrence of natural precipitation events (Fig. 2A).

While δ^2^H values peaked in the xylem five days after label application, no label was detectable in the tree crowns on the same date (Fig. 3E). Although we did not sample on a daily resolution, we found the first increase in δ^2^H values (–6.4 ± 4.3 ‰) in the tree crown on August 2, which was 12 days later than in the xylem. On August 15, δ^2^H values decreased again to pre-label values (Fig. 3E). Contrarily, we could not detect clearly labelled stemflow water in the xylem water of surrounding, unlabeled mature *F. sylvatica* and *P. abies* trees (Fig. 3D). δ^2^H values of xylem water in *P. abies* in mixed stands were similar to control trees with some minor increases on August 10 and August 26. This is supported by the absence of increased δ^2^H values in water extracted from the trees’ crowns throughout the measurement period (Fig. 3F). However, by investigating dual isotope plots of the xylem water of mature *F. sylvatica* and *P. abies* (Fig. S2), it is evident that at least some trees of both species showed an increased deuterium excess in xylem water, i.e. values plotting above the local meteoric water line (LMWL), which was potentially caused by labelled stemflow water.

In contrast, unlabeled juvenile *F. sylvatica* trees in the understory showed a clear increase in δ^2^H values in the crown water from –39.6 ± 1.1 ‰ (pre-label) to –14.2 ‰ ± 16.0 on July 24 (Fig. 3F) with the highest value detected at +585.6 ‰ (Fig. 3F). On July 31, i.e. 11 days after stemflow labelling, δ^2^H values decreased to pre-label values. As indicated by the standard error, δ^2^H values in the tree crown of juvenile *F. sylvatica* varied substantially (Fig. 3F).

### Spatial extent of stemflow

In the soil water close to the labelled trees, the highest δ^2^H value with a maximum of 1,745.8 ± 827.3 ‰ was detected on the evening of the labelling event (Fig. 3A). Four days later (July 24), when the spatial distribution of δ^2^H values was analysed, we found a strongly negative relationship between δ^2^H values in soil water and distance to a labelled tree, indicating a rather localized infiltration of stemflow (Fig. 4A). Already on the next sampling date (August 2), this relationship vanished, potentially due to frequent precipitation events (Fig. 1A).

**Figure 4:**
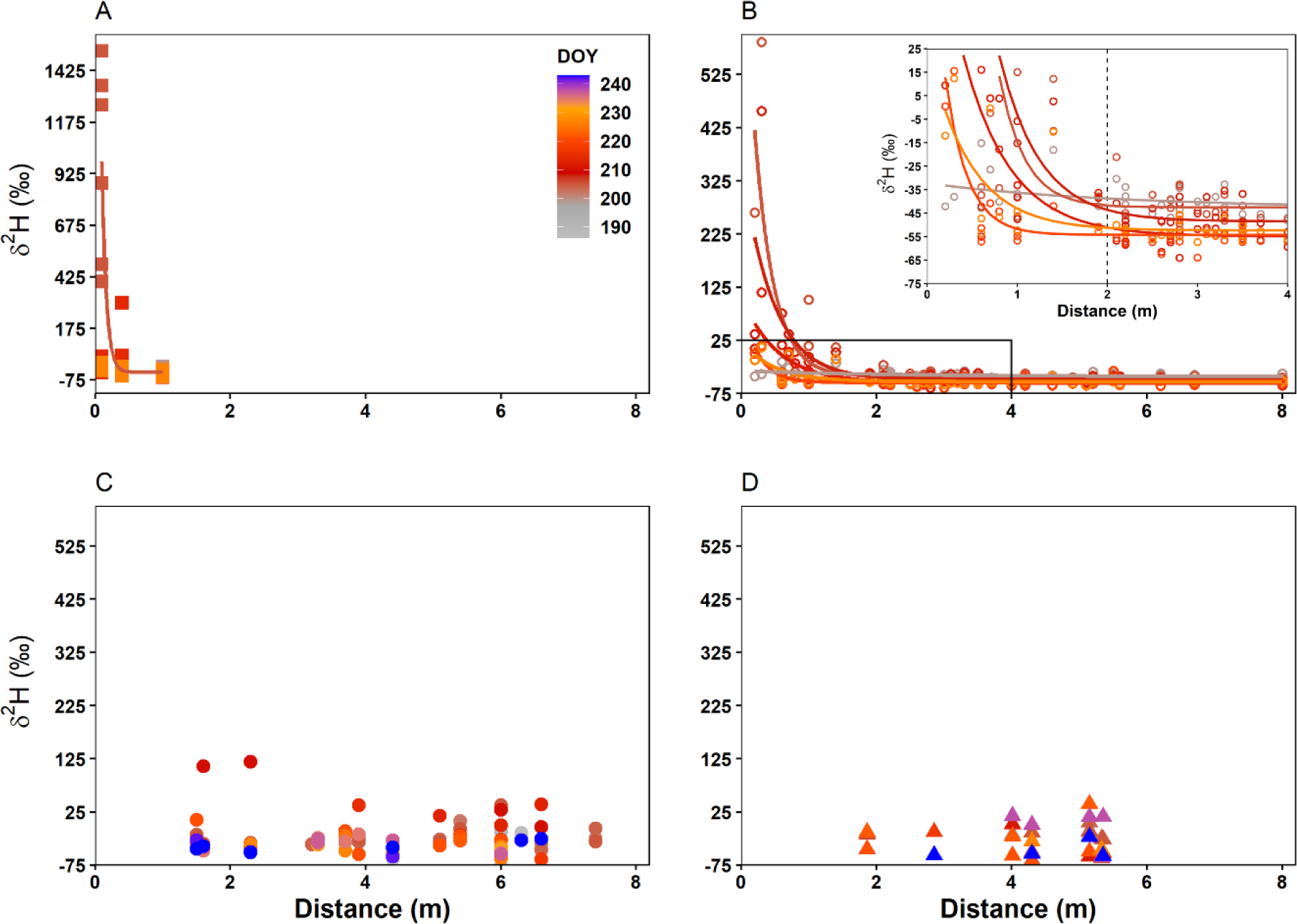
Relationship between δ^2^H values in soil water (A), crown water of juvenile *F. sylvatica* trees with zoom in (B), xylem water of mature *F. sylvatica* trees (C) and xylem water of mature *P. abies* trees (D) and distance to the closest labelled *F. sylvatica* tree. Please note the different y-axis in the panels. Regressions were modelled for each sampling day with the function *drm* in the R package *drc* (Ritz *et al*., 2015). Labelling was conducted on DOY 200, thus grey colours indicate δ^2^H values before the labelling, red colours δ^2^H values close to the labelling and blue colours δ^2^H values at the end of the measurement period. Please note the different y-scale in panel A.

Contrarily, juvenile *F. sylvatica* trees growing within a distance of 2 m around the labelled trees showed (strong) label uptake throughout the study period. The non-linear relationship of δ^2^H values in the xylem of juvenile trees and distance to the labelled tree clearly indicated a decreasing significance of stemflow with distance (Fig. 4B), explaining the large variability of δ^2^H values (Fig. 3F). With increasing time to the labelling event, the regression between δ^2^H values and distance showed lower intercepts (Fig. 4B), which is in accordance with decreasing δ^2^H values in Fig. 3F.

For mature, non-labelled *F. sylvatica* and *P. abies* trees, no relationship between δ^2^H and distance to label trees was found (Fig. 4C,D). This is supported by the comparison of δ^2^H values before and after the labelling event, which were similar. Nevertheless, two exceptions with slightly higher δ^2^H values were found in *F. sylvatica* trees at a distance of ∼2 m to a labelled tree (Fig. 4C, D).

### Stemflow contribution to xylem water

We used the *MixSIAR* model to estimate the contribution of stemflow label to water fluxes of mature *F. sylvatica* trees, which were labelled, and surrounding juvenile *F. sylvatica* trees. The model results indicated high dynamics in source water contribution to tree water uptake, with clear differences between mature and juvenile trees over the course of the experiment (Fig. 5). Isotopic compositions of xylem measured at the stem base of labelled trees revealed that 3.9 – 14.0 % of the total xylem water passing through the trunk base originated from labelled stemflow throughout the measurement period (Fig. 5A). Stemflow label was detectable at lower proportions in the stem water until the end of the measurement period in September (Fig. 4A). Measurements in the tree crown showed a slightly lower, but comparable proportion of stemflow label water uptake (7.2 – 10.1%, Fig. 4B). In general, mature *F. sylvatica* trees took up the majority of soil water from 0.20-0.40 m depth (Fig. 5A,B).

**Figure 5:**
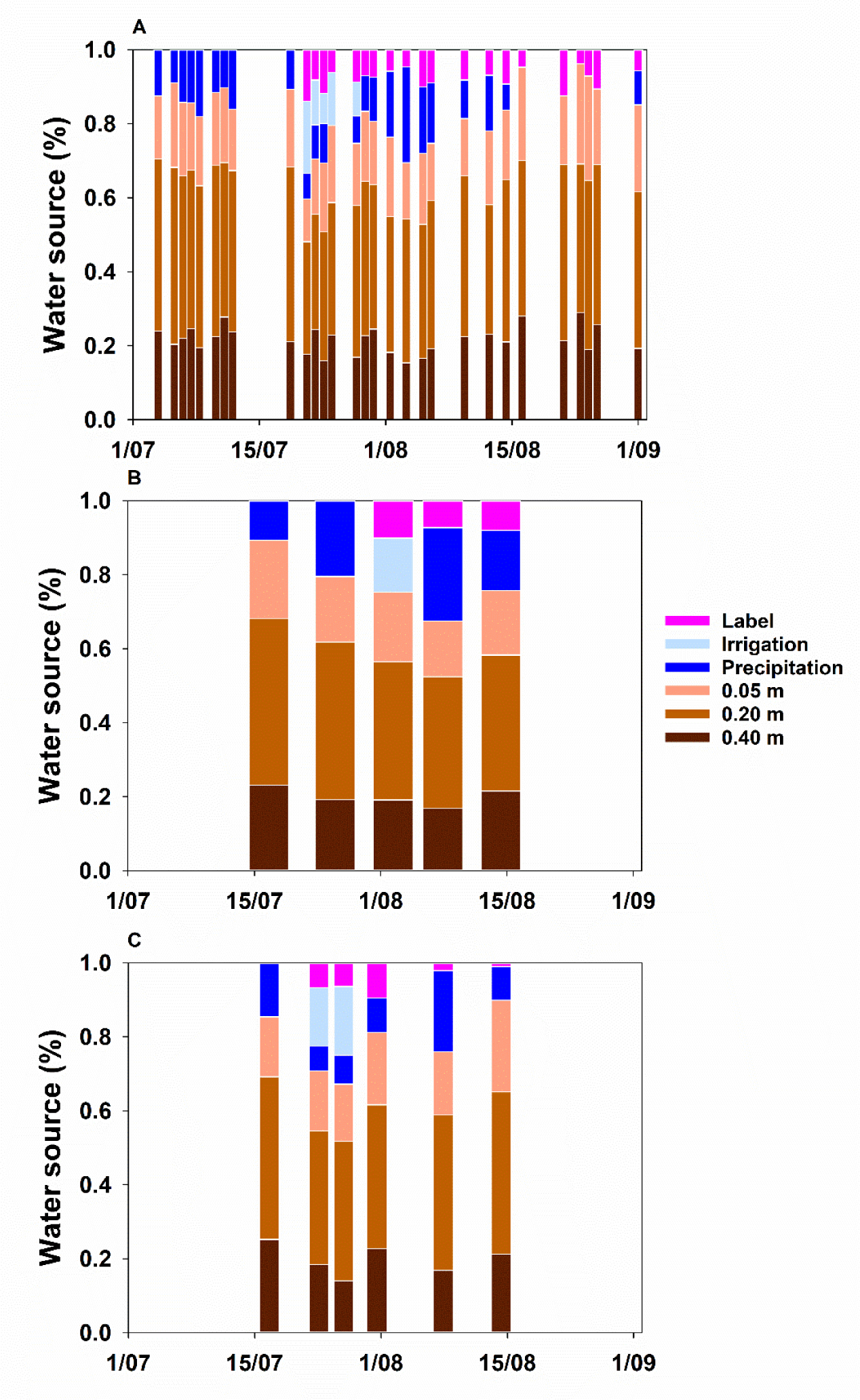
Contribution of different water sources to xylem water of labelled *F. sylvatica* trees at the stem base (n = 3-11 per day) (A), to xylem water of labelled *F. sylvatica* trees in the tree crown (n = 18) (B), and to xylem water of juvenile *F. sylvatica* trees in the tree crown (n = 45). Standard deviations are given in Table S2.

For juvenile *F. sylvatica* trees, the proportion of labelled stemflow in the xylem water was highest 11 days after the label application on July 31 (9.5 ± 12.1%, Fig 4C). Afterwards, it declined to 2.1 ± 3.5% on Aug 8 and 1.1 ± 2.4% on Aug 15, indicating a rather short transit time. Similarly to mature trees, water uptake was highest from 0.20-0.40 m depth throughout the study period. Potentially, we did not capture the maximum of stemflow label contribution in juvenile *F. sylvatica* trees, due to the lower temporal resolution compared to automated measurements at the stem base. Additionally, it has to be noted that the variability of label contribution for juvenile trees was high. Trees at a distance of > 2m to mature labelled trees showed no label uptake at all (Fig. 4B).

### Significance of stemflow for water fluxes of adult *F. sylvatica* trees

The absolute contribution of stemflow to xylem water of mature *F. sylvatica* trees ranged between 0.2 – 5.3 L day^−1^. Already two days after the label application, when sap flow rates were still high (Fig. 2C), the contribution of stemflow label was at 5.2 ± 5.9 L day^−1^ (Fig. 6). Afterwards, there was a clear decrease in absolute amounts, which is in accordance with decreasing sap flow rates. Interestingly, the amount of labelled stemflow in sap flow of *F. sylvatica* increased 4 weeks after the label application to max. values of 5.3 ± 7.1 L day^−1^. These secondary peaks coincided with (strong) precipitation events. It must be denoted that both datasets (absolute sap flow amounts and *MixSIAR* results) have rather high uncertainties.

**Figure 6:**
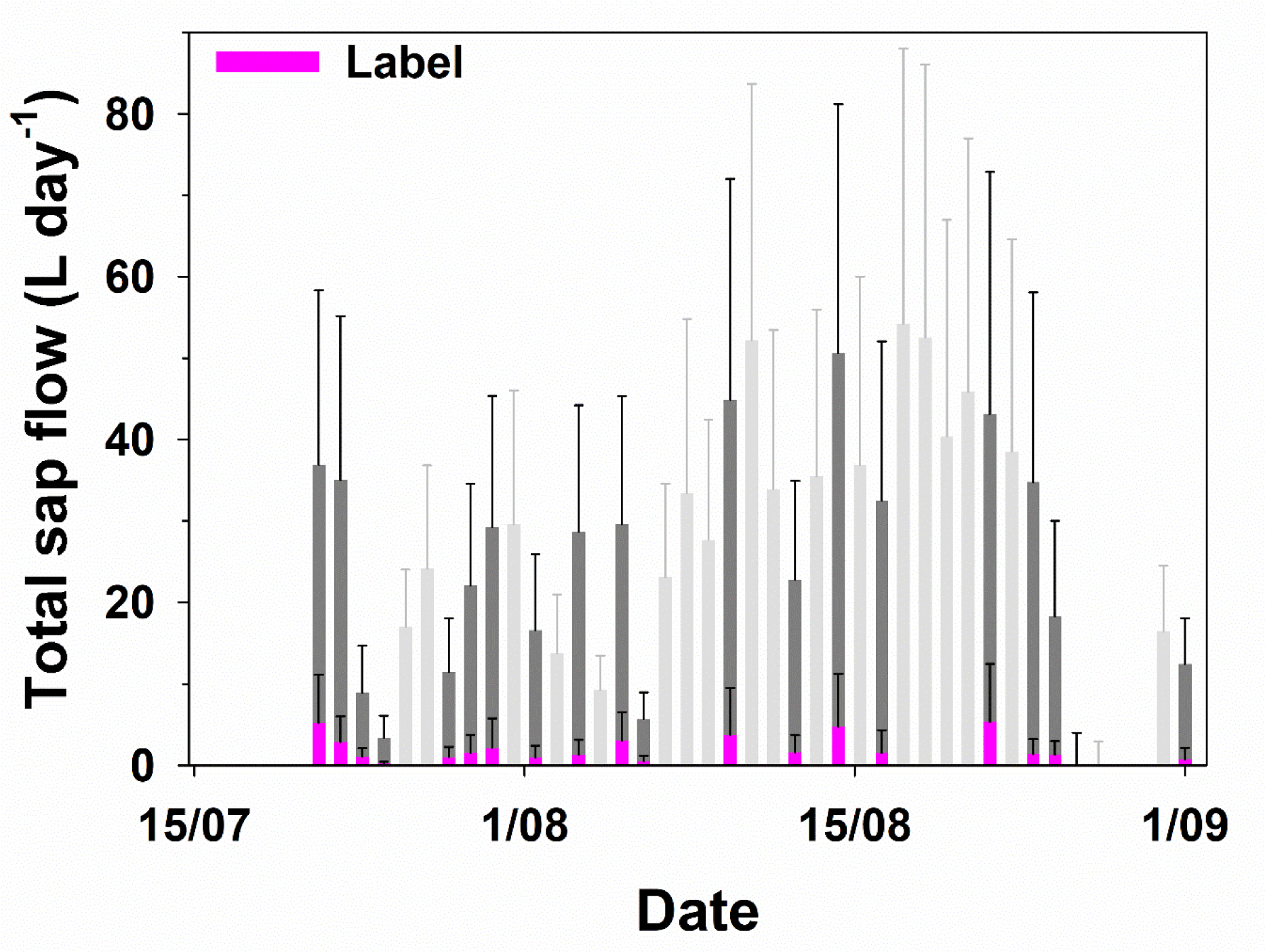
Daily sap flow of labelled *F. sylvatica* (n = 18) ± 1 SD and absolute label contribution to sap flow ± 1 SD (calculated with error propagation). Dark grey bars indicate days, where isotopic data was available and absolute label contribution could be calculated (cf. Fig. 5A). All other days are displayed in light grey.

## Discussion

With this study we demonstrate that stemflow of adult *F. sylvatica* trees provides a substantial water source for the trees themselves, but also for surrounding juvenile conspecific trees. These conspecific trees showed a high uptake of labelled water directly after label application, indicating active root water uptake from the stemflow infiltration zone of adult *F. sylvatica* trees. Although stemflow infiltrates locally at the stem base, its spatial distribution in the vegetation is larger than previously thought.

### Stemflow is a readily available and persistent water source

We could demonstrate that stemflow is an important water source for trees, which was taken up directly after precipitation events. Recent studies have shown that trees can use fresh precipitation water rapidly (Volkmann *et al*., 2016; Brinkmann *et al*., 2018; Kinzinger *et al*., 2024; Snyder *et al*., 2024; Kinzinger *et al*., 2025) – partly due to their high proportion of fine roots in the top soil (Cermák *et al*., 1993). Reported stemflow infiltration depths into the soil range from 0.07 m up to 1.22 m, depending on the precipitation amount (Návar, 2011; Spencer & van Meerveld, 2016; Zuecco *et al*., 2025). Thus, stemflow is likely to infiltrate directly into the active rooting zone of the respective trees, most likely as sub-soil preferential flow along roots, as postulated by Johnson & Lehmann (2006). This is also supported by the absence of labelled water at a distance of >0.40 m to the labelled trees in our study, indicating the presence of local stemflow infiltration hotspots (Hemr *et al*., 2023). The shares of stemflow contributing to plant water fluxes calculated in this study also agreed well with results of Kinzinger *et al*. (2024) (1-15%), who used natural abundance water stable isotope measurements on the same site. In general, results of the *MixSIAR* model for other water sources than stemflow are in line with other studies, indicating 0.20-0.40 m as important water uptake depth for *F. sylvatica* (Gessler *et al*., 2022; Kinzinger *et al*., 2024), confirming our modelling results.

While the stemflow share in xylem water was highest at the stem base of mature *F. sylvatica* trees directly after label application, we were able to detect labelled water for ∼12 days, indicating a rather long residence time compared to 1-3 days found by Snyder *et al*. (2024) in a semi-arid ecosystem. This shows that stemflow is plant available for a long period in the soil and potentially mixes with other water resources, such as fresh precipitation (Brinkmann *et al*., 2018). Accordingly, Kinzinger *et al*. (2025) recently showed that summer precipitation events were available to plants >60 days. Alternatively, trees potentially (re-)fill stem water reserves (*capacitance*) with fresh water (Kühnhammer *et al*., 2023), which might be used at a later time (Kinzinger *et al*., 2025). These processes might also explain the secondary peaks in δ^2^H enrichment and label contribution to tree water fluxes between mid-August and beginning of September, which fell into a period of frequent precipitation events. This indicates that stemflow as a water resource might also be of high relevance in times of drought and potentially provides a competitive advantage compared to other species without significant stemflow, such as *P. abies* (Jost *et al*., 2004). This is stressed by the daily contribution of stemflow to sap flow of *F. sylvatica*, which can reach 5.3 ± 7.1 L day^−1^. Thus, stemflow might play a role in the drought recovery for *F. sylvatica* (cf. Gessler *et al*., 2022), potentially allowing for faster recovery times.

With these results, we confirmed our first hypothesis that *F. sylvatica* trees benefit from their own stemflow for a rather long period, a process recently questioned by van Stan & Pinos (2024) and Snyder *et al*. (2024). However, the impact of stemflow is not confined to the base of the stem or the rooting zone of the respective tree.

### The spatial impact of stemflow in forested ecosystem extends beyond pure infiltration processes

In the soil, we could not detect any label at distances >0.40 m from the stem of a labelled tree, neither in destructive samples, nor in the continuous soil water isotope measurement system at the centre of the plots (> 1.00 m from labelled trees). These observations are consistent with other studies (Pinos *et al*., 2023), which found the infiltration area of stemflow to be < 1m² (Carlyle-Moses *et al*., 2020; Llorens *et al*., 2022), without significant surface runoff (Zuecco *et al*., 2025). Infiltration processes of stemflow are not uniform but primarily concentrated along the surface of large roots and macropores (Johnson & Lehmann, 2006). From there, the spatial movement belowground is characterized by two ways: (a) matrix flow and (b) preferential flow along roots and soil pores. Matrix flow was found to be spatially limited to the immediate vicinity of the stem. Limited lateral flow was rather observed in shallow soil layers (Schwärzel *et al*., 2012; Llorens *et al*., 2022). Yet, we found labelled water in the xylem of juvenile trees growing at a distance of up to 2 m to the labelled *F. sylvatica* trees. Van Stan & Pinos (2024) argued that the dense network of roots in an undisturbed forest can lead to intense competition for water resources, such as stemflow. *F. sylvatica* trees form dense root networks in the upper soil layers (0-0.2/0.3 m depth) (Schmid & Kazda, 2001) and root systems of different individuals often overlap. Fine roots in particular form a closely interwoven network in the topsoil and for a single tree can reach many meters laterally (Rewald & Leuschner, 2009; Lang *et al*., 2010), depending on the size and DBH of the tree (Schenk, 2006). Considering that *F. sylvatica* is one of the most competitive species belowground in temperate European forests (Bolte & Villanueva, 2006), it is likely that both mature and juvenile trees strongly profit from stemflow water. The importance of stemflow water inputs apparently decreases with increasing distance, which might be related to the decreasing fine root density with increasing distance to the tree (Rewald & Leuschner, 2009), limiting the access to infiltrated stemflow water at distances > 2 m for *F. sylvatica*. Not being able to generate large stemflow by themselves under the dense *F. sylvatica* canopy of mature trees, which can intercept >20% of precipitation (Staelens *et al*., 2008), juvenile trees might actually profit from water inputs by stemflow of mature trees. For juvenile trees, growing in close proximity to mature *F. sylvatica* trees, stemflow inputs might even provide a competitive advantage under progressing climate change, where hotter and drier conditions have been observed (Werner *et al*., 2025) and are expected to increase (for Central Europe) (Bevacqua *et al*., 2022; Luca & Donat, 2023).

In contrast, we could not detect a clear pattern in mature, unlabelled *F. sylvatica* and *P. abies* trees, growing in close proximity to labelled trees. While there was an indication that some trees took up stemflow water at certain time points, these results rather indicate that these mature trees do not rely on or profit strongly from stemflow of other trees. Mature *F. sylvatica* trees are capable of generating their own stemflow (Levia *et al*., 2010), which potentially makes the use of stemflow of other trees obsolete. While there is no significant stemflow generation by *P. abies* (Jost *et al*., 2004) due to its bark roughness, *P. abies* trees might not root deep enough (most fine roots are located in the first 0.10 - 0.25 m; (Schmid & Kazda, 2002) to reach hot spots of stemflow infiltration along preferential flow paths. Especially in mixture with *F. sylvatica*, *P. abies* shifts its root water uptake depth to even shallower layers (Hackmann *et al*., 2025). Additionally, we could not detect any labelled water in the crowns of unlabelled, mature trees during the study period. Thus, it is unclear if sporadic uptake of labelled stemflow water at the stem base by mature *F. sylvatica* and *P. abies* can be considered a relevant ecohydrological flux.

## Conclusion

The observed spatio-temporal dynamics strongly indicate that stemflow is a significant and ecohydrologically important water source for mature trees that generate it. We demonstrate that a significant proportion of stemflow was used for water uptake under non-drought conditions within 5 – 6 weeks after the event, accounting for 0.2 – 5.3 L of sap flow per day. Additionally, the impact of the stemflow label was not confined to the labelled trees, but also juvenile *F. sylvatica* could profit from these water inputs. Hence, the spatial impact of stemflow is larger than can be explained by local infiltration processes alone, most likely due to the formation of dense rooting networks and intraspecific competition for water resources. Thus, we argue that stemflow, as an important ecohydrological flux, should be considered in ecohydrological balances, modelling or experimental designs, as such neglections might lead to falsified or biased conclusions.

## Acknowledgements

We acknowledge funding from the DFG in the projects WE2681/12-1, OR480/2-1, and the CRC1537 (Project ID: 459819582). We sincerely thank Stefan Seeger, Florenz König, Britta Kattenstroth and Jonas Schwarz for technical support of the field site and support during measurement campaigns. We thank Barabara Herbstritt for assistance in analysing water stable isotope samples. Special thanks go to the City of Ettenheim for providing access to their forest.

## Competing interests

None declared.

## Author contributions

LK, JM, MW, NO, CW and SH designed the study, SK, LK, KK, CW and SH performed the experiment, SK and LK collected the data, SK, LK, JM and SH analysed the data, SK and SH interpreted the data, SK and SH wrote the manuscript with inputs from all authors. All authors critically reviewed the manuscript.

## Data Availability

The data that support the findings of this study are available from the corresponding author upon reasonable request.

## Supporting Information

***Fig. S1.*** Stemflow application system used for *Fagus sylvatica* trees.

***Table S1.*** Water stable isotope standards used in this study.

***Fig. S2.*** Dual isotope plots of measured δ^2^H and δ^18^O values in xylem water.

***Table S2A.*** Contribution of different water sources to xylem water of labelled *F. sylvatica* trees in the stem.

***Table S2B.*** Contribution of different water sources to xylem water of labelled *F. sylvatica* trees in the tree crown.

***Table S2C.*** Contribution of different water sources to xylem water of juvenile *F. sylvatica* trees in the tree crown.

## Notes

### Competing Interest Statement

The authors have declared no competing interest.

