## Supporting Information for "Water stable isotopes reveal the ecohydrological importance of stemflow for mature and juvenile European beech"

The following Supporting Information is available for this article:

**Fig. S1** Stemflow application system used for *Fagus sylvatica* trees.

**Table S1** Water stable isotope standards used in this study.

**Fig. S2** Dual isotope plots of measured $\delta^2$H and $\delta^{18}$O values in xylem water.

**Table S2A** Contribution of different water sources to xylem water of labelled *F. sylvatica* trees in the stem.

**Table S2B** Contribution of different water sources to xylem water of labelled *F. sylvatica* trees in the tree crown.

**Table S2C** Contribution of different water sources to xylem water of juvenile *F. sylvatica* trees in the tree crown.

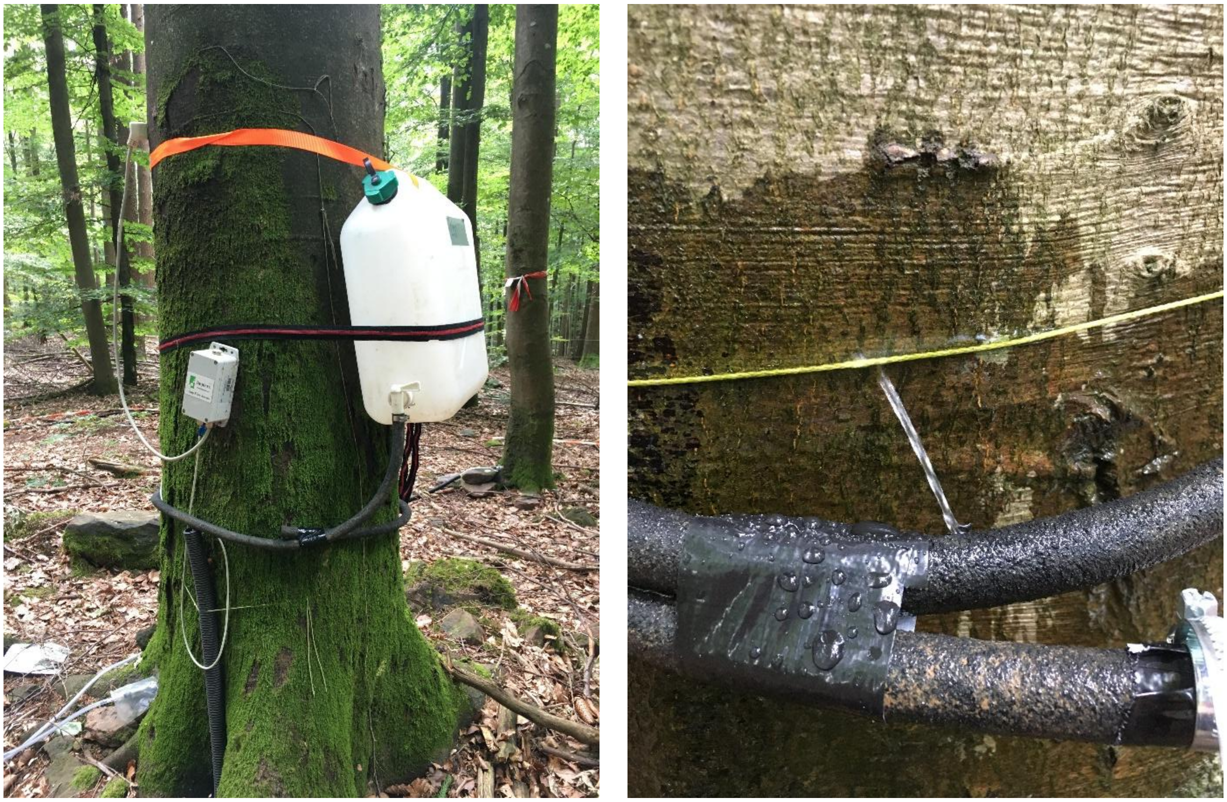

**Fig. S1** Stemflow application system used for *Fagus sylvatica* trees. $^{2}H_2O$ was filled in the white containers and was slowly applied through small holes in tubes, which were placed around the stem.

| Samples | Isotope | Standard 1 (‰) | Standard 2 (‰) | Standard 3 (‰) | Standard 4 (‰) |
|---|---|---|---|---|---|
| **Stem base (continuous measurement system)** | $\delta^2H$ | −96.12 ± 0.82 | −60.40 ± 0.78 | −0.62 ± 0.88 | NA |
| | $\delta^{18}O$ | −24.44 ± 0.29 | −17.98 ± 0.29 | −14.80 ± 0.29 | NA |
| **Tree crown (destructive sampling)** | $\delta^2H$ | −102.56 ± 0.26 | −63.79 ± 0.07 | −10.34 ± 0.05 | 54.23 ± 0.15 |
| | $\delta^{18}O$ | −25.09 ± 0.09 | −9.25 ± 0.04 | −5.20 ± 0.02 | −0.39 ± 0.03 |
| **Precipitation samples** | $\delta^2H$ | −107.96 ± 0.60 | −66.07 ± 0.60 | 1.53 ± 0.60 | NA |
| | $\delta^{18}O$ | −14.86 ± 0.16 | −9.47 ± 0.16 | 0.30 ± 0.16 | NA |

**Table S1** Water stable isotope standards used in this study.

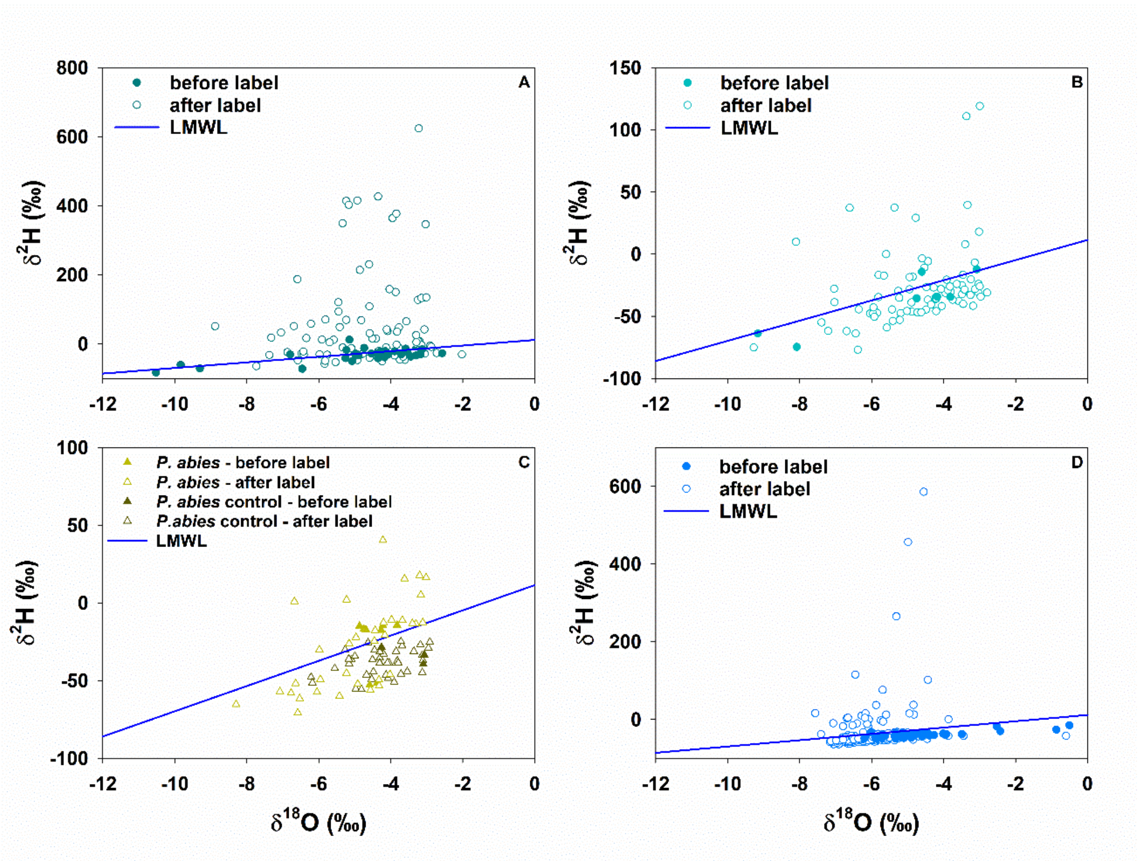


**Fig. S2** Dual isotope plots of measured $\delta^2$H and $\delta^{18}$O values in xylem water of labelled *F. sylvatica* trees at the stem base (n = 3-11 per day) (A), of unlabelled *F. sylvatica* (n = 3-9 per day) (B) and *P. abies* trees (n = 2-5 per day), including control trees for *P. abies* (n = 3-6 per day), at the stem base (C) juvenile *F. sylvatica* trees in the tree crown (n = 45) (D) with local meteoric water line (LMWL). Data is separated in points before and after the stemflow labelling event.

| Date | Measurement position | Source | Contribution + 1 SD |
|---|---|---|---|
| 4 Jul 2023 | Stem | 0.05 m | 0.17 ± 0.17 |
| | | 0.20 m | 0.47 ± 0.21 |
| | | 0.40 m | 0.24 ± 0.19 |
| | | Precipitation | 0.13 ± 0.15 |
| 6 Jul 2023 | Stem | 0.05 m | 0.23 ± 0.19 |
| | | 0.20 m | 0.48 ± 0.24 |
| | | 0.40 m | 0.20 ± 0.17 |
| | | Precipitation | 0.09 ± 0.12 |
| 7 Jul 2023 | Stem | 0.05 m | 0.20 ± 0.18 |
| | | 0.20 m | 0.44 ± 0.22 |
| | | 0.40 m | 0.22 ± 0.16 |
| | | Precipitation | 0.14 ± 0.17 |
| 8 Jul 2023 | Stem | 0.05 m | 0.18 ± 0.16 |
| | | 0.20 m | 0.43 ± 0.21 |
| | | 0.40 m | 0.25 ± 0.18 |
| | | Precipitation | 0.14 ± 0.14 |
| 9 Jul 2023 | Stem | 0.05 m | 0.19 ± 0.16 |
| | | 0.20 m | 0.44 ± 0.22 |
| | | 0.40 m | 0.20 ± 0.17 |
| | | Precipitation | 0.18 ± 0.16 |
| 11 July 2023 | Stem | 0.05 m | 0.20 ± 0.19 |
| | | 0.20 m | 0.46 ± 0.24 |
| | | 0.40 m | 0.23 ± 0.21 |
| | | Precipitation | 0.12 ± 0.16 |
| 12 Jul 2023 | Stem | 0.05 m | 0.20 ± 0.17 |
| | | 0.20 m | 0.42 ± 0.21 |
| | | 0.40 m | 0.28 ± 0.18 |
| | | Precipitation | 0.10 ± 0.13 |
| 13 Jul 2023 | Stem | 0.05 m | 0.17 ± 0.17 |
| | | 0.20 m | 0.44 ± 0.24 |
| | | 0.40 m | 0.24 ± 0.21 |
| | | Precipitation | 0.16 ± 0.17 |
| 20 Jul 2023 | Stem | 0.05 m | 0.21 ± 0.17 |
| | | 0.20 m | 0.47 ± 0.22 |
| | | 0.40 m | 0.21 ± 0.17 |
| | | Precipitation | 0.11 ± 0.13 |
| 22 Jul 2023 | Stem | 0.05 m | 0.12 ± 0.08 |
| | | 0.20 m | 0.30 ± 0.14 |
| | | 0.40 m | 0.18 ± 0.13 |
| | | Precipitation | 0.07 ± 0.10 |
| | | Irrigation | 0.20 ± 0.14 |
| | | Stemflow (label) | 0.14 ± 0.14 |
| 23 Jul 2023 | Stem | 0.05 m | 0.15 ± 0.11 |
| | | 0.20 m | 0.31 ± 0.17 |
| | | 0.40 m | 0.24 ± 0.15 |
| | | Precipitation | 0.09 ± 0.12 |
| | | Irrigation | 0.12 ± 0.11 |
| | | Stemflow (label) | 0.08 ± 0.08 |
| 24 Jul 2023 | Stem | 0.05 m | 0.19 ± 0.14 |
| | | 0.20 m | 0.35 ± 0.18 |
| | | 0.40 m | 0.16 ± 0.14 |

| Date | Measurement position | Source | Contribution + 1 SD |
| --- | --- | --- | --- |
| | | Precipitation | 0.11 ± 0.13 |
| | | Irrigation | 0.08 ± 0.09 |
| | | Stemflow (label) | 0.12 ± 0.08 |
| 25 Jul 2023 | Stem | 0.05 m | 0.21 ± 0.16 |
| | | 0.20 m | 0.36 ± 0.18 |
| | | 0.40 m | 0.23 ± 0.17 |
| | | Precipitation | 0.00 ± 0.00 |
| | | Irrigation | 0.14 ± 0.13 |
| | | Stemflow (label) | 0.06 ± 0.05 |
| 28 Jul 2023 | Stem | 0.05 m | 0.17 ± 0.13 |
| | | 0.20 m | 0.41 ± 0.18 |
| | | 0.40 m | 0.17 ± 0.14 |
| | | Precipitation | 0.07 ± 0.10 |
| | | Irrigation | 0.09 ± 0.11 |
| | | Stemflow (label) | 0.09 ± 0.10 |
| 29 Jul 2023 | Stem | 0.05 m | 0.19 ± 0.15 |
| | | 0.20 m | 0.42 ± 0.20 |
| | | 0.40 m | 0.23 ± 0.16 |
| | | Precipitation | 0.10 ± 0.12 |
| | | Stemflow (label) | 0.07 ± 0.09 |
| 30 Jul 2023 | Stem | 0.05 m | 0.17 ± 0.17 |
| | | 0.20 m | 0.39 ± 0.20 |
| | | 0.40 m | 0.25 ± 0.20 |
| | | Precipitation | 0.12 ± 0.15 |
| | | Stemflow (label) | 0.07 ± 0.12 |
| 1 Aug 2023 | Stem | 0.05 m | 0.22 ± 0.15 |
| | | 0.20 m | 0.37 ± 0.20 |
| | | 0.40 m | 0.18 ± 0.15 |
| | | Precipitation | 0.18 ± 0.11 |
| | | Stemflow (label) | 0.06 ± 0.08 |
| 3 Aug 2023 | Stem | 0.05 m | 0.15 ± 0.12 |
| | | 0.20 m | 0.39 ± 0.18 |
| | | 0.40 m | 0.15 ± 0.11 |
| | | Precipitation | 0.26 ± 0.11 |
| | | Stemflow (label) | 0.05 ± 0.06 |
| 5 Aug 2023 | Stem | 0.05 m | 0.19 ± 0.15 |
| | | 0.20 m | 0.36 ± 0.17 |
| | | 0.40 m | 0.17 ± 0.15 |
| | | Precipitation | 0.18 ± 0.12 |
| | | Stemflow (label) | 0.10 ± 0.10 |
| 6 Aug 2023 | Stem | 0.05 m | 0.15 ± 0.11 |
| | | 0.20 m | 0.40 ± 0.17 |
| | | 0.40 m | 0.19 ± 0.14 |
| | | Precipitation | 0.16 ± 0.11 |
| | | Stemflow (label) | 0.09 ± 0.11 |
| 10 Aug 2023 | Stem | 0.05 m | 0.16 ± 0.14 |
| | | 0.20 m | 0.44 ± 0.19 |
| | | 0.40 m | 0.22 ± 0.17 |
| | | Precipitation | 0.10 ± 0.12 |
| | | Stemflow (label) | 0.08 ± 0.12 |
| 13 Aug 2023 | Stem | 0.05 m | 0.20 ± 0.14 |

| Date | Measurement position | Source | Contribution + 1 SD |
|---|---|---|---|
| | | 0.20 m | 0.35 ± 0.17 |
| | | 0.40 m | 0.23 ± 0.15 |
| | | Precipitation | 0.15 ± 0.13 |
| | | Stemflow (label) | 0.07 ± 0.09 |
| 15 Aug 2023 | Stem | 0.05 m | 0.19 ± 0.16 |
| | | 0.20 m | 0.44 ± 0.20 |
| | | 0.40 m | 0.21 ± 0.17 |
| | | Precipitation | 0.07 ± 0.10 |
| | | Stemflow (label) | 0.09 ± 0.12 |
| 17 Aug 2023 | Stem | 0.05 m | 0.25 ± 0.18 |
| | | 0.20 m | 0.42 ± 0.20 |
| | | 0.40 m | 0.28 ± 0.19 |
| | | Stemflow (label) | 0.05 ± 0.08 |
| 22 Aug 2023 | Stem | 0.05 m | 0.19 ± 0.17 |
| | | 0.20 m | 0.48 ± 0.19 |
| | | 0.40 m | 0.21 ± 0.17 |
| | | Stemflow (label) | 0.12 ± 0.14 |
| 24 Aug 2023 | Stem | 0.05 m | 0.27 ± 0.19 |
| | | 0.20 m | 0.40 ± 0.21 |
| | | 0.40 m | 0.29 ± 0.18 |
| | | Stemflow (label) | 0.04 ± 0.05 |
| 25 Aug 2023 | Stem | 0.05 m | 0.28 ± 0.21 |
| | | 0.20 m | 0.46 ± 0.21 |
| | | 0.40 m | 0.19 ± 0.18 |
| | | Stemflow (label) | 0.07 ± 0.08 |
| 26 Aug 2023 | Stem | 0.05 m | 0.21 ± 0.15 |
| | | 0.20 m | 0.43 ± 0.22 |
| | | 0.40 m | 0.26 ± 0.20 |
| | | Stemflow (label) | 0.11 ± 0.12 |
| 31 Aug 2023 | Stem | 0.05 m | 0.24 ± 0.20 |
| | | 0.20 m | 0.42 ± 0.21 |
| | | 0.40 m | 0.19 ± 0.18 |
| | | Precipitation | 0.06 ± 0.11 |
| | | Stemflow (label) | 0.09 ± 0.15 |

**Table S2A** Contribution of different water sources to xylem water of labelled *F. sylvatica* trees (n = 3-11 per day) in the stem ±1SD

| Date | Measurement position | Source | Contribution + 1 SD |
|---|---|---|---|
| 18 Jul 2023 | Crown | 0.05 m | 0.21 ± 0.18 |
| | | 0.20 m | 0.45 ± 0.22 |
| | | 0.40 m | 0.23 ± 0.19 |
| | | Precipitation | 0.11 ± 0.17 |
| 26 Jul 2023 | Crown | 0.05 m | 0.18 ± 0.16 |
| | | 0.20 m | 0.43 ± 0.20 |
| | | 0.40 m | 0.19 ± 0.16 |
| | | Precipitation | 0.21 ± 0.14 |
| 2 Aug 2023 | Crown | 0.05 m | 0.19 ± 0.15 |
| | | 0.20 m | 0.37 ± 0.20 |
| | | 0.40 m | 0.19 ± 0.16 |
| | | Irrigation | 0.15 ± 0.13 |
| | | Stemflow (label) | 0.10 ± 0.12 |
| 8 Aug 2023 | Crown | 0.05 m | 0.15 ± 0.13 |
| | | 0.20 m | 0.36 ± 0.17 |
| | | 0.40 m | 0.17 ± 0.14 |
| | | Precipitation | 0.25 ± 0.16 |
| | | Stemflow (label) | 0.07 ± 0.09 |
| 15 Aug 2023 | Crown | 0.05 m | 0.17 ± 0.16 |
| | | 0.20 m | 0.37 ± 0.20 |
| | | 0.40 m | 0.22 ± 0.16 |
| | | Precipitation | 0.16 ± 0.17 |
| | | Stemflow (label) | 0.08 ± 0.10 |

**Table S2B** Contribution of different water sources to xylem water of labelled *F. sylvatica* trees (n =18) in the tree crown ±1SD.

| Date | Measurement position | Source | Contribution + 1 SD |
|---|---|---|---|
| 18 Jul 2023 | Crown | 0.05 m | 0.16 ± 0.16 |
| | | 0.20 m | 0.44 ± 0.22 |
| | | 0.40 m | 0.25 ± 0.18 |
| | | Precipitation | 0.15 ± 0.16 |
| 24 Jul 2023 | Crown | 0.05 m | 0.16 ± 0.13 |
| | | 0.20 m | 0.36 ± 0.17 |
| | | 0.40 m | 0.18 ± 0.16 |
| | | Precipitation | 0.07 ± 0.09 |
| | | Irrigation | 0.16 ± 0.14 |
| | | Stemflow (label) | 0.07 ± 0.09 |
| 27 Jul 2023 | Crown | 0.05 m | 0.16 ± 0.14 |
| | | 0.20 m | 0.38 ± 0.19 |
| | | 0.40 m | 0.14 ± 0.12 |
| | | Precipitation | 0.08 ± 0.10 |
| | | Irrigation | 0.19 ± 0.16 |
| | | Stemflow (label) | 0.07 ± 0.09 |
| 31 Jul 2023 | Crown | 0.05 m | 0.20 ± 0.17 |
| | | 0.20 m | 0.39 ± 0.20 |
| | | 0.40 m | 0.23 ± 0.17 |
| | | Precipitation | 0.10 ± 0.12 |
| | | Stemflow (label) | 0.09 ± 0.12 |
| 8 Aug 2023 | Crown | 0.05 m | 0.17 ± 0.15 |
| | | 0.20 m | 0.42 ± 0.20 |
| | | 0.40 m | 0.17 ± 0.15 |
| | | Precipitation | 0.22 ± 0.14 |
| | | Stemflow (label) | 0.02 ± 0.04 |
| 15 Aug 2023 | Crown | 0.05 m | 0.25 ± 0.20 |
| | | 0.20 m | 0.44 ± 0.22 |
| | | 0.40 m | 0.21 ± 0.18 |
| | | Precipitation | 0.09 ± 0.13 |
| | | Stemflow (label) | 0.01 ± 0.02 |

**Table S2C** Contribution of different water sources to xylem water of juvenile *F. sylvatica* trees (n =45) in the tree crown ±1SD.
